# *Scoary2:* Rapid association of phenotypic multi-omics data with microbial pan-genomes

**DOI:** 10.1101/2023.04.19.537353

**Authors:** Thomas Roder, Grégory Pimentel, Pascal Fuchsmann, Mireille Tena Stern, Ueli von Ah, Guy Vergères, Stephan Peischl, Ola Brynildsrud, Rémy Bruggmann, Cornelia Bär

**Affiliations:** Interfaculty Bioinformatics Unit and Swiss Institute of Bioinformatics and Graduate School for Cellular and Biomedical Sciences, University of Bern, 3012 Bern, Switzerland; Methods development and analytics, Agroscope, Schwarzenburgstrasse 161, CH-3003 Bern, Switzerland; Food microbial systems, Agroscope, Schwarzenburgstrasse 161, CH-3003 Bern, Switzerland; Norwegian Institute of Public Health, Oslo and Norwegian University of Life Science, Ås, Norway

**Keywords:** prokaryote, bacteria, pan-genome, metabolite, microbial genome-wide association studies, GWAS, BGWA, genotype-phenotype association, fermented food, omics

## Abstract

Genomic screening of bacteria is common practice to select strains with desired properties. However, 40-60% of all bacterial genes are still unknown, making capturing the phenotype an important part of the selection process. While omics-technologies collect high-dimensional phenotypic data, it remains challenging to link this information to genomic data to elucidate the impact of specific genes on phenotype. To this end, we present Scoary2, an ultra-fast software for microbial genome-wide association studies (mGWAS), enabling integrative data exploration. As proof of concept, we explore the metabolome of 44 yogurts with different strains of *Propionibacterium freudenreichii*, discovering two genes affecting carnitine metabolism.

## Background

The emergence of large-scale whole-genome sequencing, coupled with rapid development of tools for analyzing and sharing data, presents unprecedented opportunities to understand microbial genomics, to establish connections between genetic variations and functions, both at the level of individual organisms and within complex microbial communities. In the field of fermented foods, research focuses not only on studying the common characteristics of bacteria, but also on the individual abilities of certain bacteria to produce specific components in a product. Metabolic models can be used to gain deep insights into bacterial physiology, which is one possible approach to address these questions. However, useful models are challenging to develop, particularly for interacting microbial communities such as those present in yogurt and cheese [1]. These models make strong assumptions and are inherently limited to established and curated networks of genes and metabolites. Since the function of many bacterial genes (around 40–60% [2]) is not yet known, a more straightforward approach to learn about the function and interaction of genes is to capture the bacterial phenotype and correlate it with genomic data. While this may not lead to a holistic understanding of the microbes, it allows to select bacteria appropriate for the composition of specific fermented foods. In this context, individual genes can be key for acquiring a desired characteristic.

Although it is not an easy task to capture the full abundance of substances that make up the final nutritional composition of fermented food, recent developments in omics technologies such as mass spectrometry (MS) make it possible to capture massive metabolic profiles [3].

Even though sequencing and omics technologies are advancing rapidly, linking these data to gain an understanding of functional relationships remains a major challenge. Numerous and conceptually different approaches have been developed to integrate omics datasets, with a strong focus on human genetics and disease-related tasks such as disease subtyping or biomarker prediction [4]. Among these, only a few attempt to directly link phenotypes measured by omics technologies to genes using established human genome-wide association (hGWAS) or quantitative trait loci (QTL) methods [5]. Unfortunately, due to the differences between human and microbial genomes, hGWAS methods cannot be readily applied to microbes.

Microbial genome-wide association studies (mGWAS), sometimes termed bacterial genome-wide association (BGWA), are still a new area of research with the goal of finding genetic explanations to bacterial phenotypes [6]. The reason why the well-established methods of hGWAS cannot simply be adapted lies in the plasticity of bacterial genomes. In human, the genetic diversity is very low. This is not surprising given that the generation time of humans is around 25 years [7], population bottlenecks occurred only around 70,000 years ago [8] (around 2,800 generations) and founder effects during migration further reduced diversity [9]. In contrast, bacteria commonly have reproduction times measured in minutes and can be much older: the last common ancestor of *E. coli K-12* and *E. coli O157:H7* lived about 4.5 million years ago [10]. To illustrate the difference, most of our genes still have chimpanzee orthologs, and only 0.6% of bases [11] in a typical human genome differ from the human reference genome. Meanwhile, the core_97%_ genome of 10,667 *E. coli* genomes represents only 1.96% of the total pangenome [12]. As a result, hGWAS is typically performed by aligning reads to a human reference genome and focuses almost exclusively on single nucleotide polymorphisms (SNPs), amounting to more than 99.9% of human genomic variants [11]. In mGWAS, on the other hand, researchers more often focus on gene-presence-absence, copy-number-variants, unitigs or *k*-mers. Moreover, humans reproduce sexually, and the genome is diploid. Because of recombination, genetic variants that are in proximity have a higher chance of being co-inherited, a phenomenon termed “linkage disequilibrium” that can lead to false positives in GWAS. As bacteria reproduce clonally, the entire genome is in linkage disequilibrium and population structure becomes a strong confounding factor (pseudoreplication) [13, 14]. For this reason, classical dimension-reducing techniques popular in hGWAS, such as multidimensional scaling (MDS), might not sufficiently control false positives. Finally, bacterial genomes are very diverse, with varying numbers of circular or linear DNA molecules, sometimes with plasmids or phages, and recombination and mutation rates that may vary considerably between and even within species. While bacteria do not exchange genetic material through meiosis, recombination of DNA can happen in many species through the processes of transformation, transduction or conjugation [15, 16].

A good overview of existing mGWAS software can be found in San et al. [6]. Among the tools presented, Scoary was the most-cited software (as of February 2023), undoubtedly due to its simplicity and user-friendliness. Scoary scores binary genomic features (i.e., presence/absence of orthogenes, SNPs, unitigs or *k*-mers) for associations to a binary phenotype using Fisher’s test and accounts for population structure using a post-hoc label-switching permutation test. This post-hoc permutation test is based on the pairwise comparisons algorithm [17, 18]. A major advantage of this permutation test is that users do not need to experiment with ill-informed mutation rate parameters or inform the program about population structure [19].

According to San et al. [6], many mGWAS solutions are limited in that they lack data pre-processing functionality as well as post-GWAS methods. Moreover, Scoary was not designed to handle large sample sizes and requires binning for quantitative phenotypes. In addition, the use of mGWAS was hitherto mostly limited to single phenotypes, usually to pathogenicity and to drug resistance.

In the here presented study, 182 strains belonging to 20 different (sub-)species were selected from the strain collection of Agroscope, the Swiss center of excellence for agricultural research, which comprises over 10,000 isolates of lactic acid bacteria - a valuable legacy from a century of cheese research. Over the past decade, more than 1,300 of these isolates were sequenced and the data collected in the Dialact database. Of these selected strains, 182 yogurts were produced, each by combining one strain from the strain collection with the same starter culture. The metabolomes of these yogurts were measured using liquid chromatography MS (LC-MS) and gas chromatography MS (GC-MS), the latter measuring the yogurt’s volatile metabolites (volatiles). The aim of this study was to investigate the effect of the pan-genome of the added bacterial strains on the phenotype of the yogurts.

Here we present Scoary2, a complete re-write and extension of the original Scoary software, developed to efficiently link phenotypic multi-omics data of yogurt to microbial genomes using mGWAS and enable integrative data exploration of yogurt metabolomes. Scoary2 is significantly faster and can thus be applied to more traits as well as isolates. Moreover, the pre-processing (binning) of continuous phenotypes is now integrated and the types of genomic input-data permitted are expanded. Crucial for efficient post-GWAS data exploration of large datasets, Scoary2 includes a simple frontend implemented in HTML/JavaScript that visually and interactively integrates the data as well as optional metadata describing isolates, traits and orthogenes.

## Results

### The Scoary2 software

#### Overview

Scoary2 retains all features that are already familiar to users of original Scoary [19]. As in Scoary, the two basic inputs are i) a table that describes the genotypes (orthogenes, SNPs, *k*-mers, unitigs) present in all isolates and ii) a table containing the trait(s) of the isolates. These function as explanatory and response variables, respectively. Optionally, metadata files describing the genotypes, traits, and isolates can be added, greatly facilitating the exploration of the output. Like in original Scoary, the output is a list of significant genes per trait. A manual [20] as well as a tutorial [21] detailing how to use Scoary2 is available on GitHub. Below, we describe the improvements over original Scoary.

#### Scoary2: performance enhancements

The original Scoary software only had one software dependency (SciPy [22]) and the entire software was implemented using Python-native data structures (i.e., lists and dictionaries) only. In general, Scoary2 uses the efficient NumPy [23] and pandas [24] libraries to load and process the data. Most importantly, the pairwise comparison algorithm was reimplemented, drastically reducing the number of phylogenetic tree traversals. Gene-presence-absence and trait-presence-absence data are now represented as Boolean NumPy arrays, enabling just-in-time compilation of the pairwise comparison algorithm using Numba [25]. The new implementation of this most time-consuming step is around 40x faster than original Scoary. In addition, confidence intervals in the permutation test only depend on the topology of the gene and the number of isolates with the trait. In a dataset with many traits, the same confidence intervals may be used many times. Thus, caching confidence intervals in an SQLite database [26] reduces the number of times this expensive algorithm is executed. The modular software design makes it possible to import the pairwise comparison from the Scoary2 Python module and re-use the algorithm in different programs. Another substantial speed boost comes from enabling true multiprocessing during binarization and analysis of traits using the producer/consumer software architecture pattern. Also, Scoary2 uses a just-in-time-compiled implementation of Fisher’s test (available as a standalone Python library [27, 28]) which is orders of magnitudes faster than the reference implementation in SciPy. Moreover, original Scoary is limited to analyzing datasets with less than 3,000 isolates due to Python’s recursion limit. By dynamically adjusting this limit, Scoary2 can now analyze datasets with up to 13,000 isolates.

Using equivalent settings (*permute* = 1000, *correction* I, *p-value-cutoff* = 0.1 / *n-permut* = 1000, *multiple-testing*=native:0.1), Scoary2 is about 63 times faster at analyzing 100 randomly selected traits from the dataset described in this paper (44 isolates, 9051 genes). Scoary2 can analyze a dataset with 44 isolates and 20,000 continuous traits in around 30 minutes on an Intel i7-8565U laptop (4 cores, 8 threads, 1.80-4.60 GHz), something that was not feasible with original Scoary.

#### Scoary2: software distribution

Scoary2 can be installed using the python package manager (pip) or used through an official docker container, where all dependencies are bundled, guaranteeing easy installation far into the future, thus ensuring reproducibility.

#### Binning of continuous phenotypes

The core algorithm of Scoary is based on binary genotype and phenotype data. Scoary2 is newly capable of automatically pre-processing continuous phenotypes into binary ones. For this purpose, two Scikit-learn [29] methods, *k*-means and Gaussian mixture model, are available. The former will classify all isolates as having or lacking the trait. The Gaussian mixture model seeks to fit two Gaussian distributions and calculates the probability of each isolate having or not having the trait. By default, isolates that are classified with less than 85% predicted posterior probability are ignored from further analysis. The fitting of Gaussian mixture models can fail, and the user can decide whether to skip such traits or use k-means as a backup instead. In the data exploration app, the original continuous values are used again to calculate a histogram.

#### OrthoFinder support

The name Scoary was chosen in homage to the orthology inference software Roary [30], which transformed bacterial comparative genomics in 2015 thanks to its speed and user-friendliness [31]. However, Roary does not seem to be under active development anymore and was not included in recent *Quest for Orthologs* benchmark studies [32]. Today, OrthoFinder is the most accurate ortholog inference method according to this benchmark [32, 33]. It is under continued development and is among the most used tools in the field. As input, original Scoary uses Roary’s *gene-count* table, which indicates how many genes per orthogroup each genome has. However, this makes it cumbersome to find the relevant genes of an interesting orthogroup. While Scoary2 is still compatible with the *gene-count* table, it is highly recommended to use the *gene-list* table, produced by both Roary and OrthoFinder, where cells contain a list of gene identifiers. This way, the gene names will be shown in the data exploration app.

#### Output and data exploration app

Scoary2 produces similar tables as output as original Scoary. As San et al. [6] indicated, the ability to add annotations to orthogroups would “contribute immensely” to the utility of mGWAS tools. Therefore, Scoary2 does not just allow to add metadata to orthogroups, but also to traits and isolates. In addition, Scoary2 contains a simple data exploration app for easy inspection of the results. It was built using the JavaScript libraries Bootstrap, Papa Parse, Slim Select, DataTables, Plotly and Phylocanvas [34–39] and consists of two pages.

The first page, *overview*.*html* (Figure 1), shows a dendrogram of all traits that were analyzed. The dendrogram is calculated based on how the traits split the isolates into two sets. The distance metric used a “symmetric” Jaccard index which ensures that highly correlated and highly anti-correlated traits end up close to each other in the dendrogram. The negative logarithms of the corrected *p*-value from Fisher’s test, the *p*-value from the permutation test, and the product of the two values are presented next to the dendrogram. These plots, created with SciPy and matplotlib [22, 40], can show at least 20,000 traits. When the mouse pointer hovers over a trait, the associated metadata is presented.

**Figure 1:**
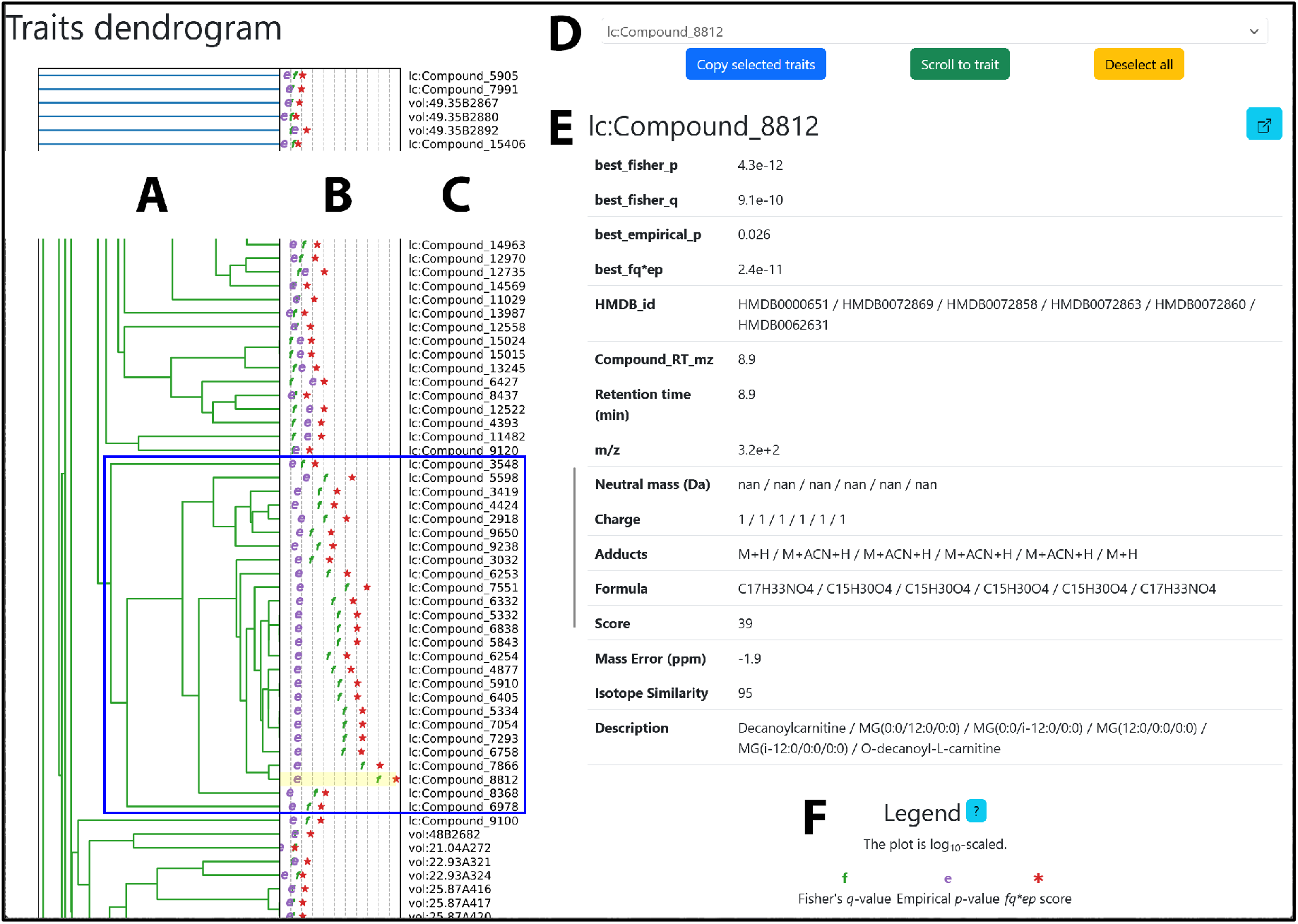
The first page (overview.html) of the Scoary2 data exploration tool. (**A**): Dendrogram of traits. The blue rectangle surrounds a cluster of carnitine-related genes. (**B**): Negative logarithms of the *p*-values calculated by Scoary2: *p*-values range from high (left) to low (right); **f** stands for the *p*-value from Fisher’s test, **e** for the *p*-value from the post-hoc test and ^**\***^ for the product of the two values. (**C**): Trait names. (**D**): Trait search and navigation tool (**E**): Trait metadata. It is updated when the mouse hovers over the traits in the dendrogram. (**F**): Plot legend.

The second page, *trait*.*html* (Figure 2), allows users to further investigate each trait. This page includes a phylogenetic tree of the isolates, where color bars indicate which isolates have the trait and which have a selected orthogroup. In addition, a pie chart shows the fraction of isolates that have the trait and how many of these have the gene. If the trait data is continuous, a histogram is also displayed. These plots are updated whenever the user clicks on an orthogroup. Below the phylogenetic tree, there are two tables. The first displays the Scoary statistics and, if present, metadata for each orthogroup. The second table is a *coverage matrix*, which shows the number of genes each isolate has from each orthogroup. If the isolates have metadata, this information is also displayed in this table. If Scoary2 uses an OrthoFinder-style *gene-list* table as input, clicks on these numbers reveal the gene identifiers. Moreover, the data exploration app can be configured to generate hyperlinks, such that clicks on gene identifiers forward the user to a certain URL, for example one where more information about the gene is available, such as its sequence and annotations. Clicks on orthogroups can also be configured to redirect to custom URLs, for example to enable a comparison of the genes.

**Figure 2:**
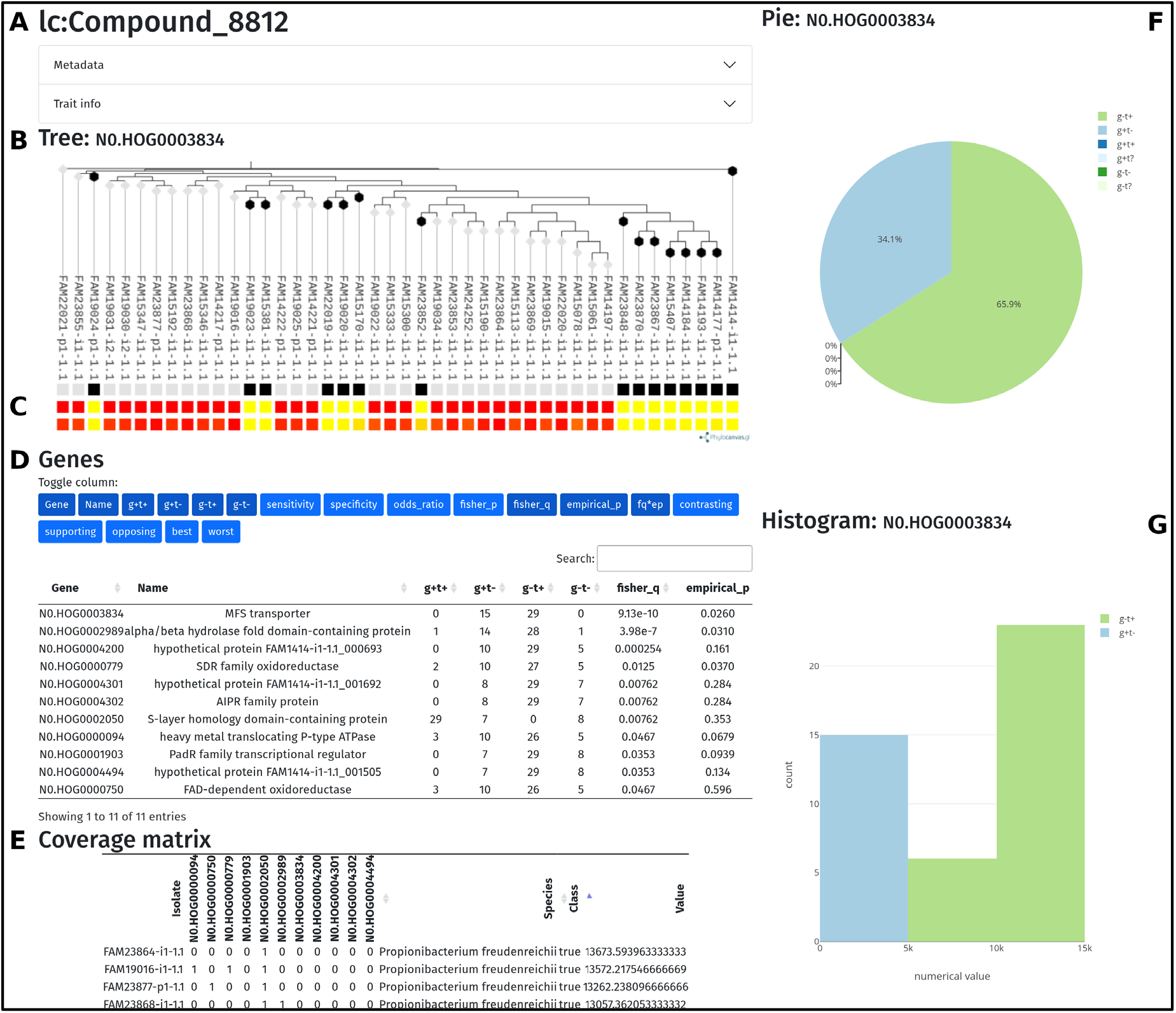
The second page (trait.html) of the Scoary2 data exploration tool. (**A**): Trait name. (**B**): Phylogenetic tree of the isolates. (**C**): Top row: presence (black) / absence (white) of orthogene. Middle row: binarized trait. Bottom row: continuous trait. (**D**): List of best candidate orthogenes with associated *p*-values. (**E**): Coverage matrix: The numbers in the cells tell the number of genes in the genome that have the annotation. (**F**): Pie chart that shows how the orthogene and the trait intersect in the dataset. (**G**): Histogram of the continuous values, colored by whether each isolate has the orthogene (g+/g-) and the trait (t+/t-).

### Scoary2 analysis of yogurt dataset

#### Dataset overview

Figure 3 A/B shows a 2D embedding of the LC-MS and GC-MS volatiles datasets that was generated using uniform manifold approximation and projection (UMAP) [41]. Notably, yogurts made with closely related strains tend to cluster together. Both datasets show one cluster dominated by yogurts made with strains from the order *Propionibacteriaceae*, and another dominated by *Lactobacillales*. Interestingly, the control yogurts which contain only the starter strains cluster with the former in the LC-MS dataset, but with the latter in the GC-MS volatiles dataset.

**Figure 3:**
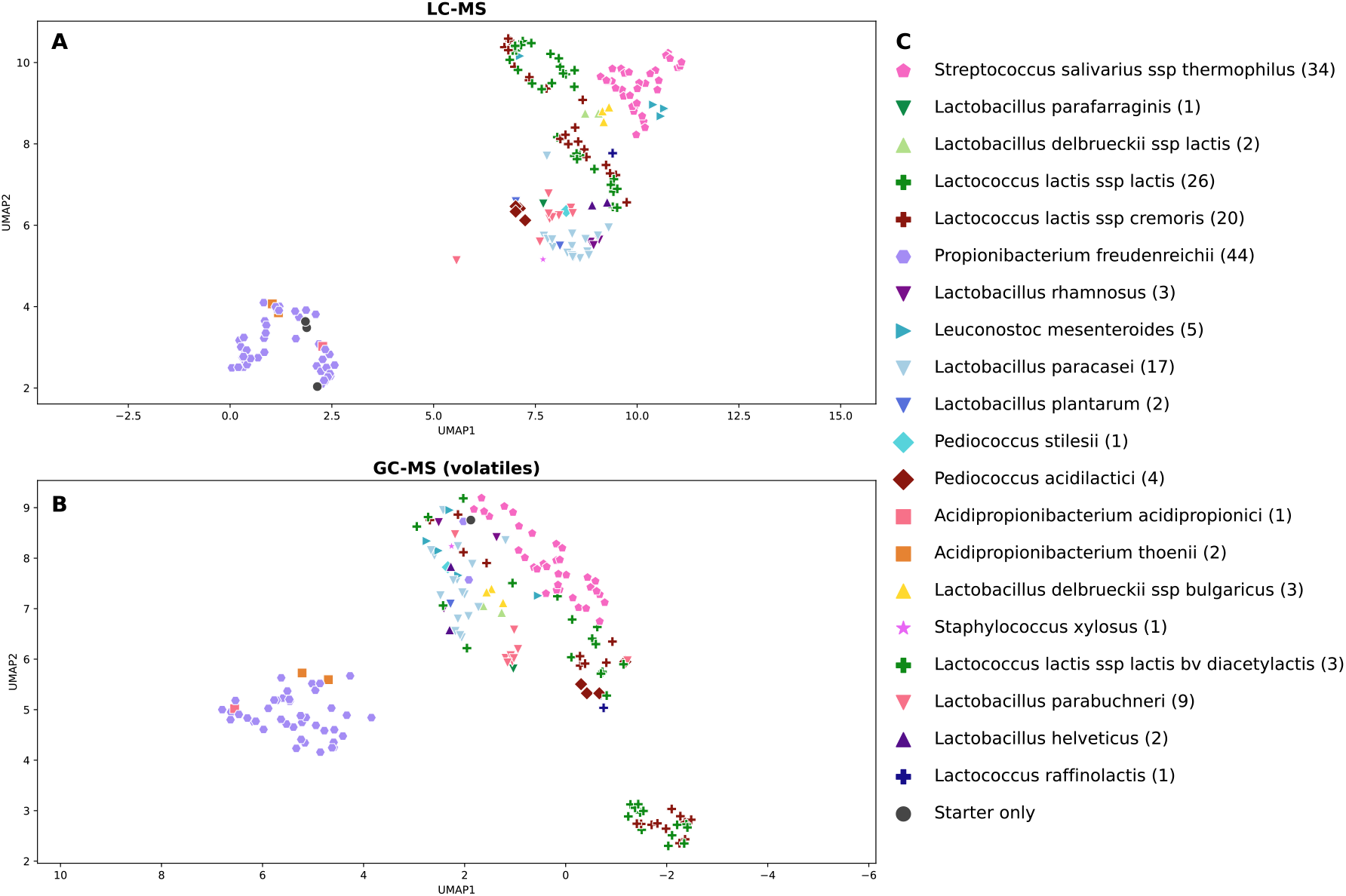
UMAP projections of mass spectrometry datasets. Each symbol represents one yogurt that was made with a different bacterial strain in addition to the starter culture YC-381. (**A**): LC-MS dataset: 2,348 metabolites. (**B**): GC-MS volatiles dataset: 1,541 metabolites. (**C**): Legend: each (sub)species has a unique combination of color and symbol. The number in brackets indicates the number of yogurts made using the respective (sub-)species.

#### Scoary2 results

With the parameters *n_cpus* = 8, *multiple_testing* = bonferroni:0.1, *n_permut* = 1000, *worst_cutoff* = 0.1, *max_genes* = 50, Scoary2 took 22 minutes to analyze the full dataset (3,889 traits, 182 isolates, 10,358 hierarchical orthogroups). As the analysis of this full dataset would go beyond the scope of this publication, as proof of concept, we show results that can be replicated by restricting the dataset to the *Propionibacterium freudenreichii* isolates. Scoary2 took only one minute to process this reduced dataset (3,889 traits, 44 isolates, 1,466 hierarchical orthogroups), with the parameters *n_cpus* = 8, *n_permut* = 1000. In comparison, original Scoary took 7.65 min to process ten traits with analogous parameters (*p-value-cutoff* = 0.1, *permute* = 1000), or approximately 50 h for the entire dataset.

The output consists of 1,256 metabolites (those with Fisher’s test *q*-values > 0.999 are automatically removed). One cluster of metabolites, highlighted with a blue rectangle in Figure 1, had particularly high *p*-values (e.g., *Compound_8812*: *q*-value Fisher’s test: 9.1 × 10^−10^, *p*-value from post-hoc test: 0.026). Because similar metabolites are clustered together (including the anti-correlated metabolites like *Compound_3032*) and each metabolite’s metadata is available in *overview*.*html*, we noticed straightforwardly that the MS database found hits with molecules with carnitine in their names for 14 out of 21 of the metabolites in this cluster (Figure 4 D). Looking at the results in more detail using *trait*.*html*, shown in Figure 2, we found that two genes correlate strongly with these metabolites: an MFS transporter and an α/β-hydrolase fold domain-containing protein. A closer look at the gene identifiers suggests that the two genes are adjacent. Furthermore, the gene loci (Figure 4 E/F) were compared using OpenGenomeBrowser [42] via custom URLs as mentioned earlier, revealing that the two genes are indeed adjacent and located in a syntenic gene cluster, one gene away from an L-carnitine CoA transferase (*caiA*). In the isolates which lack the two genes,many of the clusters were seemingly disjoined by transposases and other genes on the cluster were pseudogenized (Figure 4 E/F). Interestingly, the cluster includes the *caiABC* and *fixABCX* genes, which are associated with the anaerobic metabolism of carnitine [43]. To discover these patterns, summarized in Figure 4, all these pieces of information needed to be integrated, highlighting the importance and convenience of the data exploration app.

**Figure 4:**
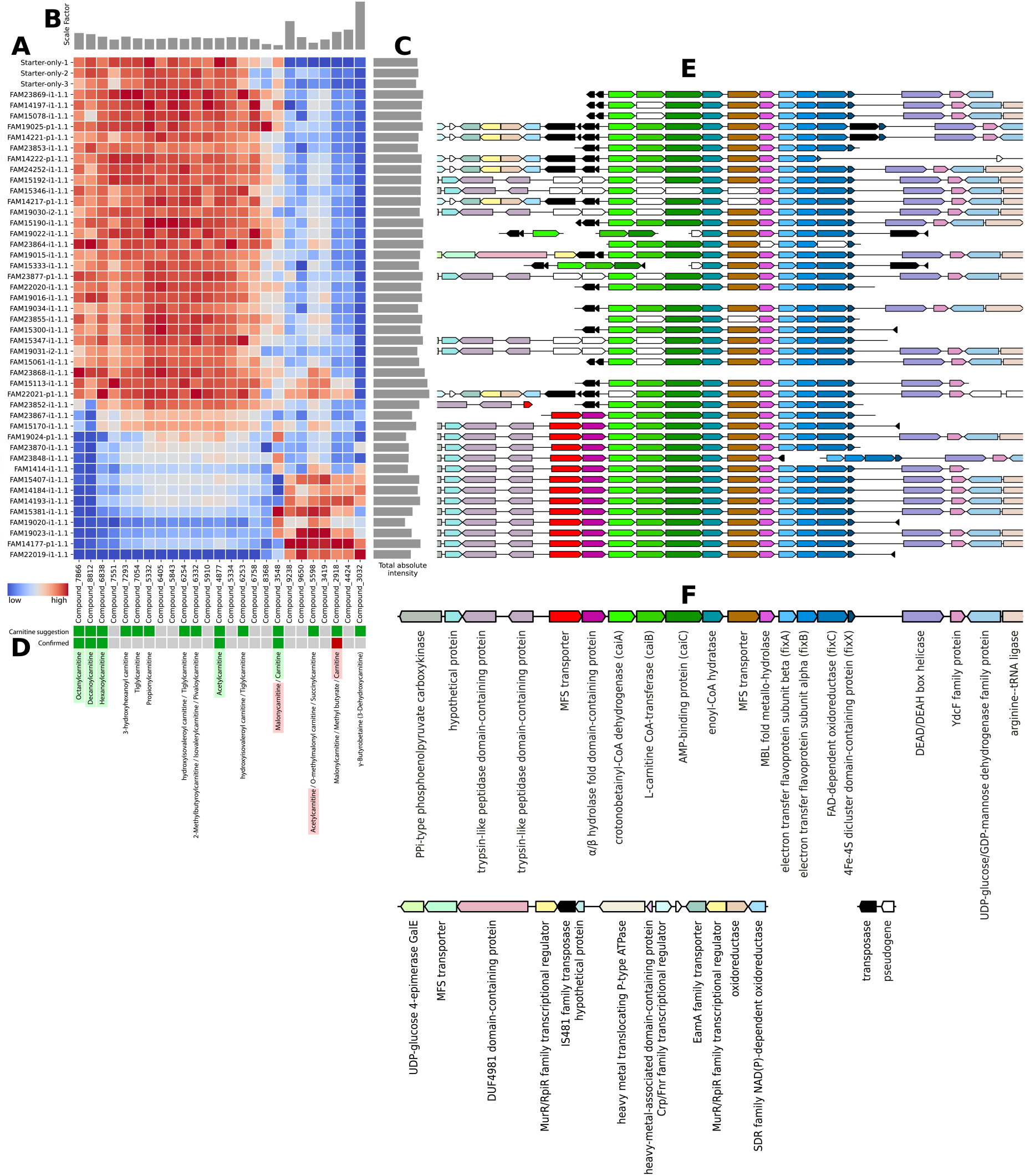
Abundance of the metabolites that correlate with the putative carnitine transporter and corresponding gene loci of three yogurts made from starter cultures only and 44 yogurts made with additional *Propionibacterium freudenreichii* isolates. (**A**): Heat map of the scaled metabolite abundances. Scale: blue (low) to average (white) to red (high). (**B**): Scale factor of each metabolite. (**C**): Sum of absolute intensities. (**D**): Color bar that indicates i) whether the mass spectrometry database suggested a match with carnitine in the name (green) or not (grey), ii) whether the suggestion could be confirmed (green) or not (red). (**E**): Comparison of the associated gene cluster spanning from the MFS transporter (red) to *fixX* (dark blue). (**F**): Annotations of the orthogenes. Orthologs are highlighted in the same color. The putative carnitine transporter highlighted in red, the *caiABC* genes in shades of green and the fixABCX genes in shades of blue.

#### Confirmation of identities for carnitine compounds

The identities of five metabolites (decanoylcarnitine, octanoylcarnitine, hexanoylcarnitine, carnitine and acetylcarnitine), assigned to the cluster detected by Scoary2 (Figure 4 D) were subsequently confirmed by LC-MS analysis of pure analytical standard solutions (Table 1).

**Table 1:** List of MFS-transporter-associated metabolites that were confirmed by standard injection.

| Metabolite | Measured m/z | Database match | CAS n° | Mass error [ppm] | Retention time error [%] |
| --- | --- | --- | --- | --- | --- |
| lc:Compound_8812 | 316.24764 | Decanoylcarnitine | 3992-45-8 | < 1 | 0.67 |
| lc:Compound_7866 | 288.21635 | Octanoylcarnitine | 25243-95-2 | 2.02 | 0.47 |
| lc:Compound_6838 | 260.18515 | Hexanoylcarnitine | 22671-29-0 | < 1 | 2.13 |
| lc:Compound_3548 | 162.11237 | Carnitine | 541-15-1 | < 1 | 8.69 |
| lc:Compound_4877 | 204.12298 | Acetylcarnitine | 3040-38-8 | < 1 | 9.66 |

Compared to yogurt made from starter cultures only, we found that two thirds of the *Propionibacterium freudenreichii* isolates did not strongly affect the composition of the carnitine-related metabolites shown in Figure 4. These yogurts are characterized by high amounts of certain acylcarnitines. In contrast, the presence in isolates of the two genes identified by Scoary2 (MFS transporter and α/β-hydrolase fold domain-containing protein), did influence the abundance of those acylcarnitines. Yogurts prepared using such isolates contain depleted amounts of acylcarnitines, particularly octanoyl- and decanoylcarnitine, and are characterized by higher amounts of carnitine, γ-butyrobetaine (putative), and certain other (putative) acylcarnitines.

## Discussion

### Translation of bacterial genomes into the metabolome of fermented foods

Although the yogurts produced in this study are multi-strain mixtures, the metabolomes of the yogurts show a clear correlation between the genetic relatedness of the added strains and the metabolomic profile of the yogurts produced. This is illustrated by the fact that the metabolomes of yogurts produced with closely related strains tend to cluster together in both MS datasets (Figure 3). Genomic differences were thus successfully translated into the metabolome of the yogurts, even though the standard manufacturing procedure does not correspond to the preferred growth conditions, such as temperature or growth time, of each species and strain. Hence, the inclusion of strains and species that differ at the genetic level in the phenotypic screening, even when done under standardized conditions, is a promising strategy to influence the composition of fermented foods. However, this taxonomy-based clustering of the metabolomic data poses a major problem when trying to find causal connections between orthogenes and metabolites, as the strongest correlations in the dataset are between the many metabolites and orthogenes that also strongly correlate with the population structure. Though these orthogenes may be good predictors of metabolism, most are not causally related to metabolites. In order to avoid spurious associations in this scenario, and to pinpoint real causal relationships, mGWAS methods such as Scoary’s pairwise comparisons are essential.

### Relevance of Carnitines in Yogurts

#### Carnitine and bacteria

L-carnitine is a ubiquitous quaternary amine compound that can be found in all kingdoms of life and is ubiquitous in many environments [44]. In bacteria, carnitine can serve multiple roles in the core metabolism, including as terminal electron acceptor and as carbon, nitrogen and energy source. Moreover, carnitine is a compatible solute and osmolyte, and is used by some bacteria as osmo- and cryoprotectant or to increase bile tolerance. Accordingly, variations in bacterial carnitine metabolism may have implications for yogurt or probiotics regarding refrigeration and gastrointestinal transit. Although many bacteria are known to be able to metabolize L-carnitine in different ways, to date, only *Sinorhizobium meliloti* is known to be able to synthesize it *de novo* [45]. In contrast, most bacteria import this molecule or direct precursors from the environment [44, 46]. Thus, transporters may be key to carnitine metabolism in bacteria. For example, the *caiT* carnitine/γ-butyrobetaine antiporter is known to be involved in anaerobic carnitine metabolism, as it imports carnitine and exports the fermentation product and is not involved in osmotic stress response. On the other hand, if a transporter is one-way only, the purpose may more likely be osmoregulation or a different metabolic route [47, 48].

#### caiABC, fixABCX, and the potential function of the identified genes

Homologs of *fixABCX* were originally characterized in *Rhizobium meliloti* where they function as a respiratory chain, providing electrons for nitrogen fixation [52]. The genes *caiABC* were first identified as part of the *E. coli caiTABCDE* operon, which is close to and co-expressed with the *fixABCX* operon and together ferment carnitine to γ-butyrobetaine in anaerobic conditions and absence of preferred substrates [43, 49, 53]. Briefly, *caiABCD* converts carnitine to γ-butyrobetaine, aided by the respiratory chain *fixABC* which provides two electrons in the reduction step (Figure 5).

**Figure 5:**
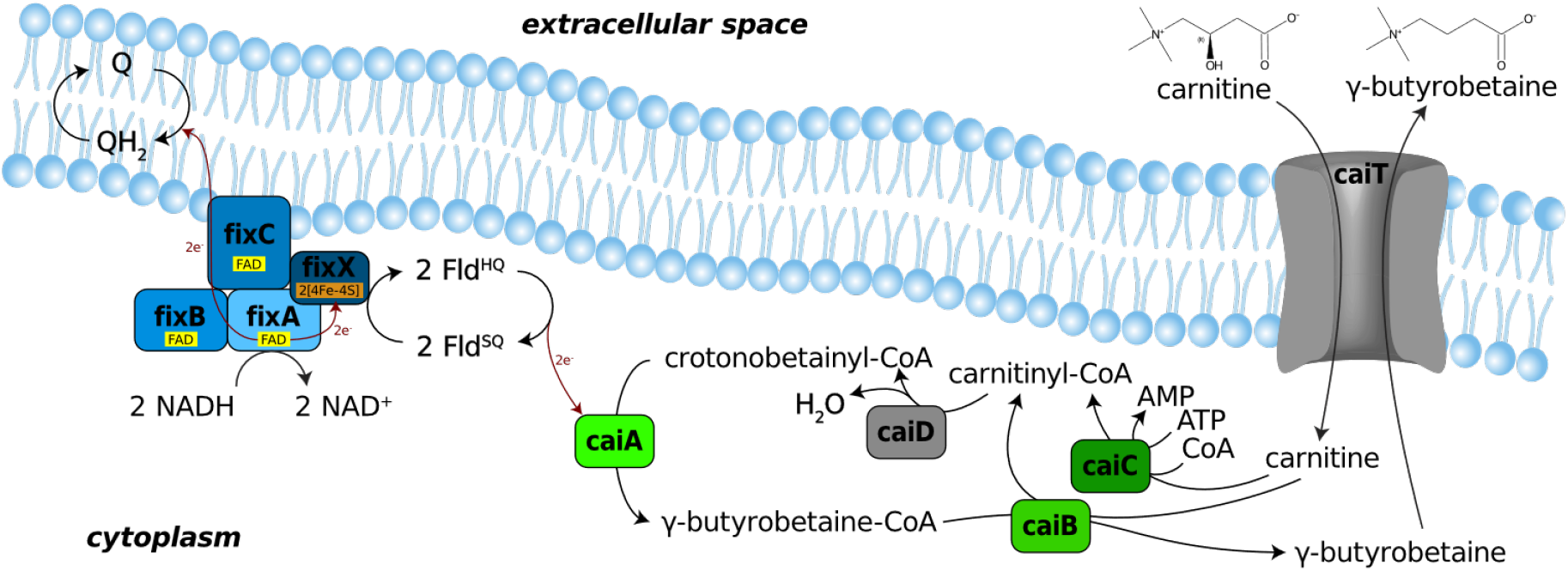
Anaerobic Carnitine Reduction in *Escherichia coli*, adapted from Walt 2002 [49], Bernal 2008 [50] and Ledbetter 2017 [51]. Proteins that were not found in any of the tested *Propionibacterium freudenreichii* strains are colored grey. ***caiT***: antiport of carnitine and γ-butyrobetaine. ***caiC***: generates initial carnitinyl-CoA. ***caiD***: dehydration of carnitinyl-CoA to crotonobetainyl-CoA. ***caiA***: reduction of crotonobetainyl-CoA to γ-butyrobetainyl-CoA using electrons from *fixABCX*. ***caiB***: recycles CoA moiety. ***fixABCX***: Oxidation of NADH is coupled to reduction of ubiquinone (Q) and flavodoxin semiquinone (Fld^SQ^), which then delivers electrons to a terminal electron acceptor. **legend**: FAD: flavin adenine dinucleotide; [4Fe-4S]: iron-sulfur cluster; Fld^SQ^: flavodoxin, semiquinone form; Fld^HQ^: flavodoxin, hydroquinone form; Q: ubiquinone; QH_2_: ubiquinol

However, the selected *Propionibacterium freudenreichii* isolates are lacking homologs of the crotonobetainyl-CoA hydratase *caiD* and the carnitine/γ-butyrobetaine antiporter *caiT*. Instead, between *caiABC* and *fixABCX*, we find an MFS transporter and an enoyl-CoA hydratase (Figure 4), which might fill these gaps in the pathway. On the other hand, the two genes identified by Scoary2 are also an MFS transporter and a hydrolase, and since only the strains with these genes have a strong impact on the carnitine composition of the yogurt (Figure 4), it appears that the full operon is required to permit efficient import of precursors and fermentation of carnitine in *P. freudenreichii*. This is supported by the apparent degradation of the gene cluster through transposases and pseudogenization in many genomes where the two genes were lost.

One piece of evidence suggests that the MFS transporter identified by Scoary2 is an acylcarnitine importer: In the *P. freundenreichii* strain FAM23848, a transposase has split *fixABC* from the rest of the operon. This appears to result in continued acylcarnitine import and deacylation, but inhibited reduction of carnitine to γ-butyrobetaine, leading to significant accumulation of carnitine (Figure 4).

If our interpretation is correct and the *P. freudenreichii* strains with the complete operon can indeed use carnitine as terminal electron acceptor, it could enable them to persist better in the human gut, where such redox reactions are a key ecological pressure [54].

#### Carnitine and humans

Even though humans can synthesize L-carnitine endogenously from the essential amino acids L-methionine and L-lysine, 75% is obtained through diet [55], which is why it has been termed a “conditionally essential nutrient”. One of the richest sources of carnitine is red meat, but bovine milk (about 169 µmol/L [56]) and milk products also contain carnitine [57]. Humans require L-carnitine to transport long- to short-chain fatty acids in and out of the mitochondrion [44]. This “carnitine shuttle” is rate-limiting for fatty acid oxidation (FAO) [58].

There are two known pathways that link carnitine to human diseases via microbiota metabolism: First, two isobaric (*m/z* = 160.133) microbe-produced structural analogs of L-carnitine (3-methyl-4-(trimethylammonio)butanoate and 4-(trimethylammonio)pentanoate) have recently been shown to hamper FAO in the mural brain by inhibiting the carnitine shuttle. Molecules with matching *m/z* have been linked to type 2 diabetes, preeclampsia, and nephropathy in type 1 diabetes. This is plausible because these diseases are associated with mitochondrial dysfunction or incomplete FAO [59]. However, no molecules with matching m/z were found in the dataset presented in this paper. Second, certain gut microbes, for example *Acinetobacter*, metabolize dietary L-carnitine to trimethylamine (TMA), which is oxidized in the liver to trimethylamine-*N*-oxide (TMAO). These metabolites are well-known risk factors of cardiovascular disease, though controversy exists as to which of these metabolites is the real culprit [60–62].

Either way, the metabolism of carnitine by the microbiota is of growing scientific interest and may be influenced by the metabolites in yogurt or the large amounts of bacteria it contains. This is indicated by two independent experiments by Burton et al. [63], where ingestion of a probiotic yogurt resulted in a lower postprandial TMAO response in urine and plasma, compared to non-fermented milk. In addition, recently discovered enzymes from the MttB superfamily of the gut bacterium *Eubacterium limosum* were found to demethylate L-carnitine and other TMA precursors and may deplete TMA/TMAO levels through precursor competition [64, 65]. In addition to impacting the postprandial TMAO response, Burton et al. also found that the ingestion of probiotic yogurt resulted in a different production of several bile acids [66], indicating that dietary fat metabolism in humans can be modulated through fermented foods via pathways involving carnitine and bile salts [63, 66].

### Potential use in microbial specialized metabolites discovery

To the best of our knowledge, Scoary2 is the first software that makes the study of large phenotypic multi-omics datasets using mGWAS feasible. This approach may constitute a novel discovery strategy for microbial metabolites, thereby providing the potential to accelerate progress in microbiology, drug discovery, and targeted production of functional fermented food to support human health [67, 68]. After all, as outlined in van der Hooft et.al. [69], traditional methods are based on established knowledge and labor-intensive experiments, such as activity-guided fractionation of metabolite extracts. These were complemented by genome and metabolome mining approaches. More recently, a “metabologenomic integration” approach was developed that combines high throughput metabolomics with genomics [69]. However, this approach does not take population structure into account and is limited to biosynthetic gene clusters (BGCs), which are challenging to predict, and depend on high quality genome sequences as well as existing knowledge [70–73]. Scoary2 on the other hand, is conceptually simpler and therefore applicable to a wider range of data, in addition to being easier to use. It is fast enough to process entire metabolomes, cannot just take BGCs but all orthogenes into account, is aware of population structure and does not rely on existing knowledge and thus represents a valid alternative in that context.

### Post-GWAS methods enable analysis of large phenotypic datasets

We strongly agree with San et al. on the immense utility of post-GWAS methods [6]. To our knowledge, Scoary2’s post-GWAS data exploration app stands out amongst other mGWAS tools, being able to integrate (i) the detected associations between traits and genes, (ii) relations between traits, (iii) relations between isolates and (iv) metadata describing traits, genes and isolates.

These innovations are very convenient for small datasets, but an absolute necessity for datasets with many traits. The dendrogram of traits in *overview*.*html* helps discover groups of (anti-)correlated traits, and the *p*-values plots help to prioritize them. The presence of trait metadata enabled us to notice quickly that many metabolites of one cluster were annotated as carnitines. Navigating to *traits*.*html* with only one click allows us to see the phylogeny of the isolates as well as the distribution of the selected trait and the highest-scoring orthogene. The orthogene annotations may also be insightful here. The gene IDs in the *coverage matrix* may reveal that certain orthogenes are often close to each other on the genome, indicating an operon. If the trait is numeric, the histogram may be useful to gauge how strongly the trait varies in the dataset and whether the data points contradicting the hypothesis might just have been incorrectly classified during binarization. If the app is connected to external comparative genomics tools, it becomes easy to study the candidate gene in more detail. In our example, OpenGenomeBrowser [42] enabled us to discover that the two genes most strongly associated with carnitines are located on the same gene cluster and near an *L-carnitine CoA transferase*, providing more evidence for a causal relationship.

Because the output of most mGWAS tools is structurally similar, i.e., consisting of coefficients for genes and traits, this app offers the possibility to be adapted to other tools or even to develop standardized output formats. Thus, a single generic data exploration app could be developed and used by many mGWAS tools.

### Comparison with existing mGWAS approaches

The field of mGWAS software is very diverse. Various conceptually different approaches have been developed and refined, and as a result, different tools require different input types and yield conceptually different outputs. The main result from LASSO and Random Forest is the model’s predictive performance. The model itself may be harder to understand, as LASSO may randomly choose one of multiple highly correlated genes and drop the others, and Random Forest does not yield correlation coefficients for the genes at all. Linear mixed models yield a straightforward *p*-value for each gene but are based on hard-to-verify assumptions about bacterial evolution. Homoplasy based methods like treeWAS [74] and Scoary give multiple *p*-values for different types of association scenarios, arguably requiring more careful interpretation. Consequently, tools based on different approaches are difficult to compare. Moreover, benchmarks are often carried out based on simulated datasets, and it is difficult to tell how closely they imitate bacterial evolution and real datasets. We noticed that Scoary and treeWAS were evaluated using simulations that emphasized the evolutionary scenarios they were designed to detect [19, 74], while the simulations from Saber et al. [14], benchmarking linear-model-based tools, did not investigate the effect of homoplastic mutations. We recommend that future research should compare the various approaches using realistic simulations and real datasets and flesh out guidelines as to which approach and tool is recommended in which scenario.

### Limitations of the Scoary2 algorithm

#### Fisher’s test

Fisher’s test is a simple and fast test that measures how strongly a gene and a trait correlate. To determine a *causal* relationship in mGWAS, however, its assumptions are violated, and the resulting *p*-values should rather be interpreted as scores. For users who simply want to learn which traits are associated with specific clusters in a tree without any assumptions on causal relation, Fisher’s test is nonetheless useful.

#### Pairwise comparisons

To be as generalizable and widely applicable as possible, the pairwise comparisons algorithm is devoid of any explicitly defined models of evolution and sacrifices some statistical power. For example, a gene whose presence is one hundred percent correlated with a particular phenotype might not be considered significant if the variant-phenotype combination is clustered on a single branch, in other words, if it can be traced back to a single event in the phylogenetic history of the input data. However, we prefer the pairwise comparisons algorithm to explicitly defined models because in our opinion, the mutation rates at every branch in the tree are most often unknown or unavailable. Thus, in Scoary2, only the branching pattern of the phylogenetic tree matters. This means that any errors in its topology could confound results.

A clear downside to the pairwise comparisons algorithm is that it can only deal with binary phenotypic events and not continuous or Brownian motion-type transitions. In Scoary2, phenotypes measured on a continuous scale are automatically binarized with either k-means or a Gaussian Mixture Model. For the former, there is a risk of improper phenotypic classification, and the latter discards values that do not clearly fit either of the gaussian means, leading to a reduced dataset to draw conclusions on. The latter issue is partially mitigated by manual inspection of the numerical values in *traits*.*html*.

#### Future directions

In the future, tests that can better exploit numerical data, can detect several types of evolutionary scenarios, or have higher statistical power could be added to Scoary2. Possible candidates are the three tests from treeWAS [74], though there is still room for the development of new algorithms [75].

## Conclusions

We expanded Scoary’s applicability to datasets containing tens of thousands of traits by significantly increasing the performance of the algorithm. Moreover, we added a novel interactive data exploration tool that combines trait, genotype, and isolate metadata, greatly facilitating the interpretation of results and crucial for timely exploration of large datasets. We illustrated Scoary2’s capabilities by applying the software to a large MS dataset of 44 yogurts made from different strains of *Propionibacterium freudenreichii*, allowing us to identify novel genes involved in carnitine metabolism. Scoary2 is, to the best of our knowledge, the first software that makes it feasible to study large phenotypic multi-omics datasets using mGWAS. It enables and facilitates the discovery of previously unknown bacterial genotype-phenotype associations and can thus help overcome a major bottleneck in microbial research, namely the unknown role of many genes and their impact on the phenotype. Therefore, it may significantly contribute to fermented food research, accelerating and facilitating the development of fermented food products with specific properties. In addition, Scoary2 has the potential for broader application, for example in basic microbial research, drug discovery and clinical research, and could thus considerably impact microbiological science in the future.

## Methods

### Yogurt production

Lactose-free, homogenized, pasteurized, semi-skimmed (1.5%) milk purchased from a local retailer was used for yogurt production (Aha! IP Suisse, Migros, Switzerland). Fermentation was carried out overnight (16 hours) at 37 °C using the yogurt culture Yoflex^®^ YC-381 (Chr. Hansen, Denmark) containing *Lactobacillus delbrueckii* subsp. *bulgaricus* and *Streptococcus thermophilus*, as well as one of the selected strains from the Liebefeld culture collection (Table S1). The yogurts were stored at -20 °C until analysis.

### GC-MS (volatiles) dataset

Untargeted volatile analysis was carried out using an Agilent 7890B gas chromatography (GC) system coupled with an Agilent 5977B mass selective detector (MSD) (Agilent Technology, Santa Clara, CA, USA). For volatile analysis, 250 mg of yogurt containing 25 µl ISTD (Paraldehyde 0.5 ppm, Tetradecane 0.25 ppm and D4-Decalactone 0.5 ppm) diluted in water were placed in 20 mL HS vials (Macherey-Nagel), hermetically sealed (blue silicone/Teflon septum (Macherey-Nagel)) and measured in a randomized order. After incubation of the samples for 10 min at 60 °C, the headspace was extracted for 5 min at 60°C under vacuum (5 mbar) as described by Fuchsmann et al., 2019 [76], using the Vacuum transfer in trap extraction method. The trap used was a Tenax TA (2/3 bottom) / Carbosieve S III (1/3 top) (BGB analytics). The temperature of the trap was fixed at 35°C and the temperature of the syringe at 100°C. The sorbent and syringe were dried for 20 min under a nitrogen stream of 220-250 mL min^− 1^. Desorption of the volatiles took place for 2 min at 300°C under a nitrogen flow of 100 mL min^−1^. For this purpose, the programmable temperature vaporization injector (PTV) was cooled at 10°C for 2 min, heated up to 250°C at a rate of 12°C sec^− 1^ and held for 20 minutes in solvent vent mode. After 2 min the purge flow to split vent was set to 100 mL min^−1^. The separation was carried out on a polar column OPTIMA FFAPplus fused silica capillary column 60m x 0.25mm x 0.5µm (Macherey-Nagel) with helium as the carrier gas at a flowrate of 1.5 mL min^-1^ (25.3 cm sec^−1^). The oven temperature was held for 5 min at 40°C, followed by heating up to 240°C at a rate of 5°C min^−1^ with a final holding time of 55 min. The trap was reconditioned after injection at a nitrogen flow of 100 mL min^− 1^ for 15 min at 300°C. The spectra were recorded in SCAN mode at a mass range between m/z 30 to m/z 350 with a gain at 10 with a solvent delay of 4 minutes. The samples were measured twice in random order. Only compounds that were detected in > 50% of QCs were retained (1,541 metabolites).

### LC-MS dataset

Untargeted metabolomic analysis was performed using an UltiMate 3000 HPLC system (Thermo Fisher Scientific) coupled to maXis 4G+ quadrupole time-of-flight mass spectrometer (MS) with electrospray interface (Bruker Daltonik GmbH). Chromatographic separation was conducted on a C18 hybrid silica column (Acquity UPLC HSS T3 1.8 µm 2.1 × 150 mm, Waters, UK), reversed phase at a flow rate of 0.4 mL min ^−1^. The mobile phase consisted in ultrafiltered water (Milli-Q^®^ IQ 7000, Merck, Germany) containing 0.1% formic acid (Fluka^(tm)^, Honeywell, USA) (A), and acetonitrile (Supelco^®^, Merck, Germany) with 0.1% formic acid (B), with the following elution gradient (A:B): 95:5 at 0 min to 5:95 at 10 min; 5:95 from 10 to 20 min; 95:5 from 20 to 30 min. The spectra were recorded from m/z 75 to m/z 1500 in positive ion mode. Detailed MS settings were as follows: collision-induced dissociation: 20 to 70 eV, electrospray voltage: 4.5 kV, endplate offset: 500 V, capillary voltage: 3400 V, nitrogen flow: 4 mL min^-1^ at 200°C, spectra acquisition rate: 1 Hz in profile mode, resolution: 80,000 FWHM. The yogurt samples were measured in triplicates in random order and the values averaged afterwards.

The QC-based robust locally estimated scatterplot smoothing signal correction method was applied for signal drift correction [77] using R (v.3.1.2) [78]. Metabolites with poor repeatability, i.e., detected in < 50% of QCs, were removed, as well as metabolites with a relative standard deviation > 30% in the QC samples. Features that had a median in the QC samples that was < 3 times higher than the median calculated for the blanks were also excluded. This reduced the number of metabolites from 17,310 to 2,348.

### Identification of carnitines

The Human Metabolome Database (27) was used with a 5-ppm mass accuracy threshold for the identification of a selection of metabolites. Identity suggestions from databases were then confirmed by MS fragmentation data (when available) and with the injection of pure standards solutions. All standards were purchased at Sigma-Aldrich (Sigma-Aldrich Chemie GmbH, Buchs, Switzerland).

### OrthoFinder

Hierarchical orthogroups were called using OrthoFinder [33] version 2.5.4 with default parameters.

### Locus plots

The gene locus plots (Figure 4) were generated using OpenGenomeBrowser [42], which utilizes DNA Features Viewer [79], and modified using Adobe Illustrator [80].

## Declarations

### Ethics approval and consent to participate

Not applicable.

### Consent for publication

Not applicable.

### Availability of data and materials

#### Software

The Scoary2 source code is publicly available at https://github.com/MrTomRod/scoary-2/ under the MIT License [81]. An official docker container (troder/scoary-2) is available on Docker Hub [82].

#### Datasets

The genomes of the 44 *Propionibacterium freudenreichii* used in this study were uploaded to NCBI GenBank and available under BioProject PRJNA946676. This data is also available via the OpenGenomeBrowser demo instance hosted at https://opengenomebrowser.bioinformatics.unibe.ch/ [83]. The combined LC-MS and GC-MS datasets and the hierarchical orthologs file generated by OrthoFinder are available in the Mendeley Data repository (doi: 10.17632/yytybr3t4y.1) under the CC BY 4.0 license [84].

### Competing interests

The authors declare that they have no competing interests.

### Funding

This research was funded by Gebert Rüf Stiftung within the program “Microbials”, grant number GRS-070/17 and the Canton of Bern.

### Authors’ contributions

GV, RB, CB, UvA, GP and TR conceived the project. GV and RB provided funding to the project. UvA and GP produced the yogurts. GP measured the LC-MS yogurt metabolome. PF and MTS measured the GC-MS volatiles yogurt metabolomes. GP identified the carnitine metabolites. SP and TR explored different ways of analyzing the data. TR programmed the Scoary2 software with advice from OB. TR, CB and GP analyzed the data. TR, CB and OB wrote the manuscript with input from all authors. TR created the figures. All authors read and approved the final manuscript.

## Acknowledgments

We thank Arthur Volant for adding Boschloo’s exact test to scipy [22] at our request, even though we ended up not using it. Furthermore, we thank Zahra Sattari for her contribution to the production of the initial yogurts.

## Notes

https://github.com/MrTomRod/scoary-2/

https://scoary.bioinformatics.unibe.ch/

https://data.mendeley.com/datasets/yytybr3t4y

### Competing Interest Statement

The authors have declared no competing interest.

